# Viral evolutionary dynamics predict Influenza-Like-Illnesses in patients

**DOI:** 10.1101/2021.01.31.429026

**Authors:** Christopher D. Wallbank, Stéphane Aris-Brosou

## Abstract

Viral infections such as those caused by the influenza virus can put a strain on healthcare systems. However, such a burden is typically difficult to predict. In order to improve such predictions, we hypothesize that the severity of epidemics can be linked to viral evolutionary dynamics. More specifically, we posit the existence of a negative association between patients’ health and the stability of coevolutionary networks at key viral proteins. To test this, we performed a thorough evolutionary analysis of influenza viruses circulating in continental US between 2010 and 2019, assessing how measures of the stability of these coevolutionary networks correlate with clinical data based on outpatient healthcare visits showing Influenza-Like Illness (ILI) symptoms. We first show evidence of a significant correlation between viral evolutionary dynamics and increased influenza activity during seasonal epidemics, and then show that these dynamics closely follow the progression of epidemics through each season, providing us with predictive power based on genetic data collected between week 20 and week 40/52, that is one to fifteen weeks prior to peak ILI. Viral evolutionary dynamics may hence be used by health authorities to further guide non-pharmaceutical interventions.

## 1 Introduction

Recent human history has been repeatedly plagued by viral outbreaks such as the 1918 H1N1 “Spanish flu” (Kilbourne, 2006), or the novel 2009 H1N1 pandemic (Smith *et al.*, 2009) – pandemics that aggravate the already heavy burden of seasonal epidemics, estimated to be as high as 650,000 deaths globally (Paget *et al.*, 2019). In any case however, these numbers are hard to predict ahead of/or as the outbreak unfolds. While standard epidemiological models can inform us on which non-pharmaceutical intervention works best to limit casualties (Davies *et al.*, 2020), and while phylodynamics can reveal the genetic structure of an epidemic (du Plessis *et al.*, 2021), models predicting the next viral strain have shown little predictive power (Sandie and Aris-Brosou, 2014), and approaches predicting burden on healthcare systems are scarce.

The case of influenza might help in this regard, as both the World Health Organization (WHO, 2020) and the Centers for Disease Control and Prevention (CDC, 2019a) closely monitor influenza activity in humans worldwide, in particular in the US. To this effect, the CDC enrolls around 3,000 healthcare providers that report numbers and percentages of patients showing symptoms of Influenza-Like-Illnesses (ILI), defined as a fever (a temperature ≥ 37.8°C) and a cough and/or sore throat, on a weekly basis, since the 1997-98 influenza season (CDC, 2019b). These weekly updates report, among others, ILI values that represent the number of patients with ILI divided by the total number of patients seen, which hence stands for a good indicator of ILI burden on the healthcare system. These data are then aggregated by state for all 50 states, plus the District of Columbia, Puerto Rico, and the US Virgin Islands, and are available since the 2010-11 season, starting right after the emergence of the pandemic 2009 H1N1 strain (CDC, 2019b). In parallel to these clinical data is a wealth of sequence data deposited in public repositories such as GenBank (Bao *et al.*, 2008), where the data can be filtered by country, season (from week 40 to week 39 of the following year), influenza subtype, host, and gene. Information about the state where each sequence was collected is available through the name of each sequence. It is therefore possible to match, in an aggregated manner, clinical and genetic data on a state by state basis, as well as on a weekly basis since 2010-11, and hence to test for the existence of genetic predictors of ILI burden. The outstanding question remains as to which genetic predictors to use for this purpose.

Previous work suggests that outbreaks and epidemics affect the evolutionary dynamics of select viruses: more specifically, outbreaks where found to be associated with a break-down of coevolving amino acid, leading to what was characterized as a destabilization of their coevolutionary networks (Aris-Brosou *et al.*, 2017). This results is sensible as experimental evidence shows that influenza viruses evolve in such a way that when some amino acid sites mutate, there may be a compensatory mutation (Gong *et al.*, 2013). When many amino acids interact in such a way, they are considered to evolve in a correlated manner, often forming networks of coevolving residues (Nshogozabahizi *et al.*, 2017), networks that are not limited to viruses as they are also found in bacteria (Dench *et al.*, 2020). These networks are thought to stabilize a genome from an evolutionary point of view (Aris-Brosou *et al.*, 2019), a stability that is compromised during an outbreak, potentially because of a lack of compensatory mutations. Because this previous work was limited to case studies, it remains unclear whether such associations can be generalized, and more critically how they translate in the clinic. To address this outstanding issue, we tested for the existence of a correlation between viral evolutionary dynamics and ILI burden, focusing on the analysis of the two main influenza antigens, the hemagglutinin and the neuraminidase genes, whose products are respectively involved in cell entry and exit of viral particles (Nelson and Holmes, 2007), found in the two subtypes commonly circulating in humans since 1968: H1N1 and H3N2. Our prediction was that if Influenza activity is indeed linked to destabilization, then the coevolving amino acid networks that are most disconnected should match to high Influenza levels in that state and season. Through a time window analysis, we show that maximum predictive power is reached when analyzing genetic data collected between week 20 and week 52, two to four weeks prior to peak ILI activity.

## 2 Materials and Methods

### 2.1 Data retrieval

Unweighted ILI data (*i.e.*, the percentage of cases of Influenza that tested positive as reported by contributing healthcare providers) were retrieved from the CDC website using the FluView portal at gis.cdc.gov/grasp/fluview/fluportaldashboard.html, for each weekly report dating from October 4, 2010, the first available date in the report and the start of the 2010-11 season, through March 10, 2020, or about two-thirds of the last season. These data were first summarized for each state and for each season by the mean ILI.

Corresponding hemagglutinin and neuraminidase coding sequences were retrieved from the NCBI Influenza database (Bao *et al.*, 2008) for the same range of dates for viruses circulating in the US. Only “full length plus” sequences, that may be missing start/stop codons, with complete collection dates, including pandemic H1N1 sequences were downloaded. Identical sequences were kept, as those could have been collected in different states. Altogether, this led us to retrieve 5,292 HA and 9,386 NA sequences for H1N1, and 16,499 HA and 14,423 NA sequences for H3N2.

### 2.2 Phylogenetic analyses

These four datasets were then aligned using Muscle ver. 3.8.31 (Edgar, 2004). Upon visual inspection, sequences that appeared misaligned were deleted, and indels caused by sequencing errors were removed. FastTree ver. 2.1.1 (Price *et al.*, 2010) was used to generate a first phylogenetic tree for each of the four datasets, assuming the General Time-Reversible +Γ model of evolution (*e.g.,* Aris-Brosou and Rodrigue, 2019). Sequences with extreme branch lengths were removed, leaving us with 5,288 HA and 9,331 NA sequences for H1N1, and 16,493 HA and 14,403 NA sequences for H3N2.

As we focused exclusively on the continental US, the “lower 48’s”, we removed all sequences collected in Alaska, Hawaii, the Virgin Islands, and the District of Columbia based on the sequence names. We corrected spelling errors, resolved city names to their state name, removed environmental sequences and sequences with odd locations that were not matching a correct state name. Remained for the final analyses 4,474 HA and 8,530 NA sequences for H1N1, and 15,197 HA and 13,147 NA sequences for H3N2. The four datasets were then split into sub-alignments by state and by season, an influenza season running from week 40 to week 39 of the following year, hence generating at most 48 × 12 = 576 state- and season-specific alignments for each gene and each subtype.

### 2.3 Reconstruction of networks of coevolving sites

For each gene and subtype, these 576 alignments were analyzed as above with FastTree to create phylogenetic trees, that were mid-point rooted with Phytools ver. 0.2.2 (Revell, 2012), and node labels (aLRT SH-like support values) were removed as they caused parsing errors at the next stage. HyPhy ver. 2.3.3 (Pond *et al.*, 2004) was then employed to infer co-evolving residues with a modified SpiderMonkey (Poon *et al.*, 2008) script (available from github.com/sarisbro). Briefly, the MG94×HKY85 codon substitution model was used to reconstruct non-synonymous substitutions in the codon sequences at each node of the tree. These reconstructions were then transformed to a binary matrix, where rows and columns represent the unique branches and amino acid positions, respectively. A Bayesian Graphical Model (BGM) was then used to find the pairs of sites that find evidence of correlated evolution. Each node represents a codon, and edges originating from a node represent a correlated relationship best explaining the data (a parent node having influence over a child node). To prevent overcomplicating the BGM, a maximum of two parents was assumed for any child. This dependence was estimated for each pair of codons by its posterior probability, itself inferred using a Markov chain Monte Carlo (MCMC) sampler that was run for 1,000,000 steps. A burn-in of 10,000 steps was assumed to be long enough to reach stationarity, and a total of 9,900 samples were evenly taken in what remained of each chain. To check for convergence, each MCMC sampler was run twice, and all downstream analyses were hence performed on each replicate, hence globally assessing the robustness of our results to sampler convergence. The two smaller temporal analyses had convergence issues and therefore were run for 10^6^ steps, with a burn-in of 5 × 10^5^ steps, and sampling a total of 95,000 samples

The igraph package ver. 1.2.4.2 (Csardi and Nepusz, 2006) was then used to create the networks of co-evolving residues, and to compute the network summary statistics: diameter (the maximum average greatest distance between any two nodes), average path length (the mean length of every potential path between any two nodes), betweenness (how often a node is included in all the shortest paths between two other nodes). *α* centrality (a measure of the influence of nodes over others), assortativity (testing whether or not nodes that are similar are likely to be connected), dyad census (a classification of the relationship for every pair of nodes), transitivity (probability that adjacent vertices are connected), and mean eccentricity (the average greatest distance between any two nodes). These network summary statistics were calculated over a series of 100 posterior probability thresholds ∈ (0.1, 0.99), representing weak to strong evidence for co-evolution.

## 3 Results and Discussion

### 3.1 Whole-season predictors of ILI

First, in order to test for the existence of a global correlation between viral evolutionary dynamics and ILI, we analyzed state-specific and season-specific sequence alignments for the HA and the NA genes from H1N1 (the pandemic strain) and H3N2 subtypes between the2010-11 and 2019-20 seasons. Note that while it can be expected that these sequences are unevenly collected through time and across states, we could not find any sequence in the database for HA in H1N1 for the last three seasons surveyed, 2017-18 to 2019-20 (Fig. S1-S2).

In all cases, these analyses included data running from week 40 of a given year to week 39 of the following year, dates that reflect the official start and the end of influenza seasons, as can be shown either by non-aggregated data (Fig. S3), or with the weekly-aggregated ILI values derived from the WHO (Fig. 1A). After alignment, these sequence were subjected to a phylogenetic reconstruction to detect pairs of (codon) sites evolving in a correlated manner based on Bayesian Graphical Models (BGMs; Poon *et al.*, 2008). Convergence of the BGMs was assessed based on the estimated posterior probabilities from two independent runs (Fig. S4). Such pairs of sites have previously been shown to form networks (Nshogozabahizi *et al.*, 2017; Dench *et al.*, 2020), that can then be analyzed based on statistics used in social network analyses such as average path length or eccentricity (Newman, 2018). Here we computed eleven such statistics, and used them first to further assess the convergence of the MCMC samplers used by the BGMs, running each analysis twice. Our results show that for all statistics, except for *α*-centrality of HA in H1N1, both runs lead to almost identical results (Fig. S5).

**Figure 1.**
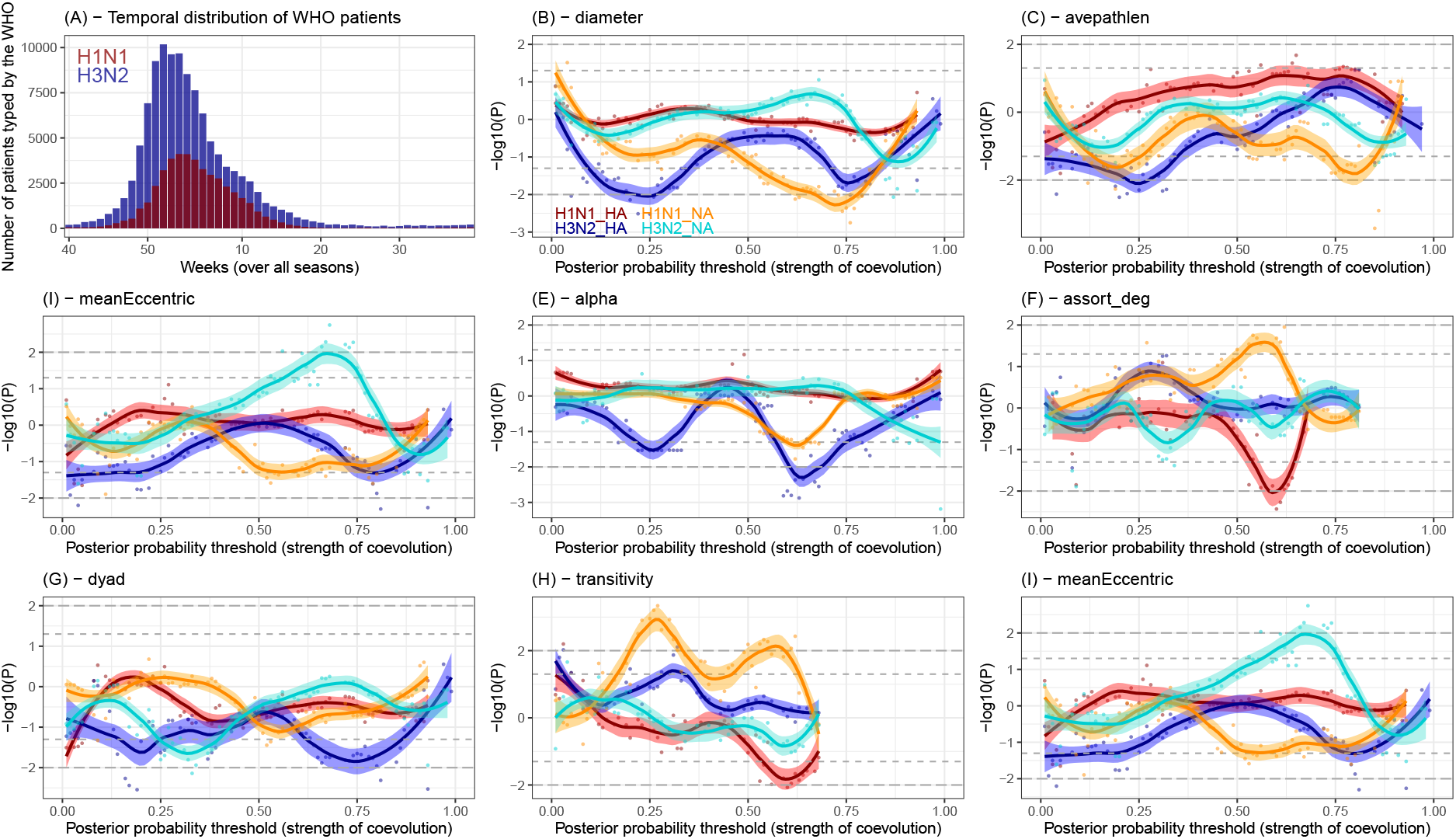
Network analyses based on genetic data from the entire season for the HA and NA genes in H1N1 and H3N2 subtypes. (A) Distribution of unweighted ILI values summed per week over the entire nine seasons, from 2010-11 to 2018-19 based on WHO data for H1N1 (dark red) and H3N2 (dark blue). The next eight panels show the significance of the robust regressions for each network statistic at a particular posterior probability threshold (the strength of coevolution) against total unweighted ILI value. Negative values indicate a negative slope, and vice-versa for positive values. Gray horizontal lines indicate significance thresholds (dash: 5%; long dash: 1%). LOWESS regressions are shown with the 95% confidence envelope for each subtype and each gene: warm colors for H1N1 (red for HA, orange for NA) and cold colors for H3N2 (blue for HA, turquoise for NA).

From previous work comparing non-pandemic and pandemic viruses, we expected that with increasing ILI, the diameter, average path length, eccentricity, and betweenness would decrease, while transitivity would increase, in particular for weakly interacting sites (around a posterior probability *PP* = 0.25, Aris-Brosou *et al.*, 2017). Our results here generally support these previous findings, in that we observe significant and negative correlations between network statistics and total ILI values for diameter (Fig. 1B), average path length (Fig. 1C), betweenness (Fig. 1D), as predicted, but also for *α*-centrality (Fig. 1E), assortative degree (Fig. 1F) and dyad (Fig. 1G). Among these, HA in H3N2 almost systematically shows a negative correlation at weak evidence for coevolution (*PP* ≈ 0.25), while NA for H1N1 shows a negative correlation at stronger evidence for coevolution (*PP* ≈ 0.75). Altogether, these negative correlations imply that amino acid residues are coevolving less as ILI increases, so that coevolutionary networks tend to dissociate. On the other hand, transitivity and mean eccentricity show positive correlations with total ILI for NA in H1N1 and H3N2, respectively (Fig. 1H-I). These statistics indicate that networks become smaller as ILI increases. Altogether, these results show that, mainly for NA in H1N1 and HA in H3N2, an increase in ILI is associated with a disruption of the coevolutionary dynamics at these two antigens, which, again, is consistent with our previous results (Aris-Brosou *et al.*, 2017).

### 3.2 Look-ahead predictors of ILI

To go beyond mere associations between network structures and ILI during an entire season, and to assess the utility of the above results in terms of public health, we split the sequence data into four different and overlapping temporal windows, beginning before the official start of each season to assess carryover effects: window 1 ran from week 20 (20 weeks before the start of the season – this is approximately when sequences deposited in GenBank start accumulating) to week 30 (10 weeks before the start of the season), window 2 ran from week 20 to week 40 (the official start of the focal season), window 3 from week 20 to week 52 (end of calendar year), and window 4 spanned weeks 20-10. We elected to start each analysis from week 20 as by then, epidemics are usually almost over (Fig. 1A), and data-deposition in GenBank reaches its nadir or lowest rate (Fig. S5). Because H3N2 is usually the subtype most impacting human populations since its emergence in 1968 (Jester *et al.*, 2020), HA and NA sequences from this subtype are about twice as abundant as for H1N1 in GenBank (Fig. S5). Furthermore, because of the missing H1N1 HA sequences for the past three seasons (Fig. S1), our data contained fewer HA than NA sequences for H1N1, when it is this latter gene sequences that are usually less sequenced (Fig. S5B). As a result, we focused the temporal analyses on H3N2 exclusively, for both HA and NA. In each case, we used genetic data coming from each time window to predict the mean ILI for the *entire* focal season.

Window 1 should serve as a control, as it starts and ends before the start of the focal season, and hence should not have accumulated enough genetic data to have any predictive value with respect to ILI (Fig. 2A). As expected, none of the network statistics showed any significance at the 1% level, except dyad census for NA, which was negatively predicting upcoming ILI at moderately interacting residues (Fig. 2B-I), and should henceforth be interpreted with care.

**Figure 2.**
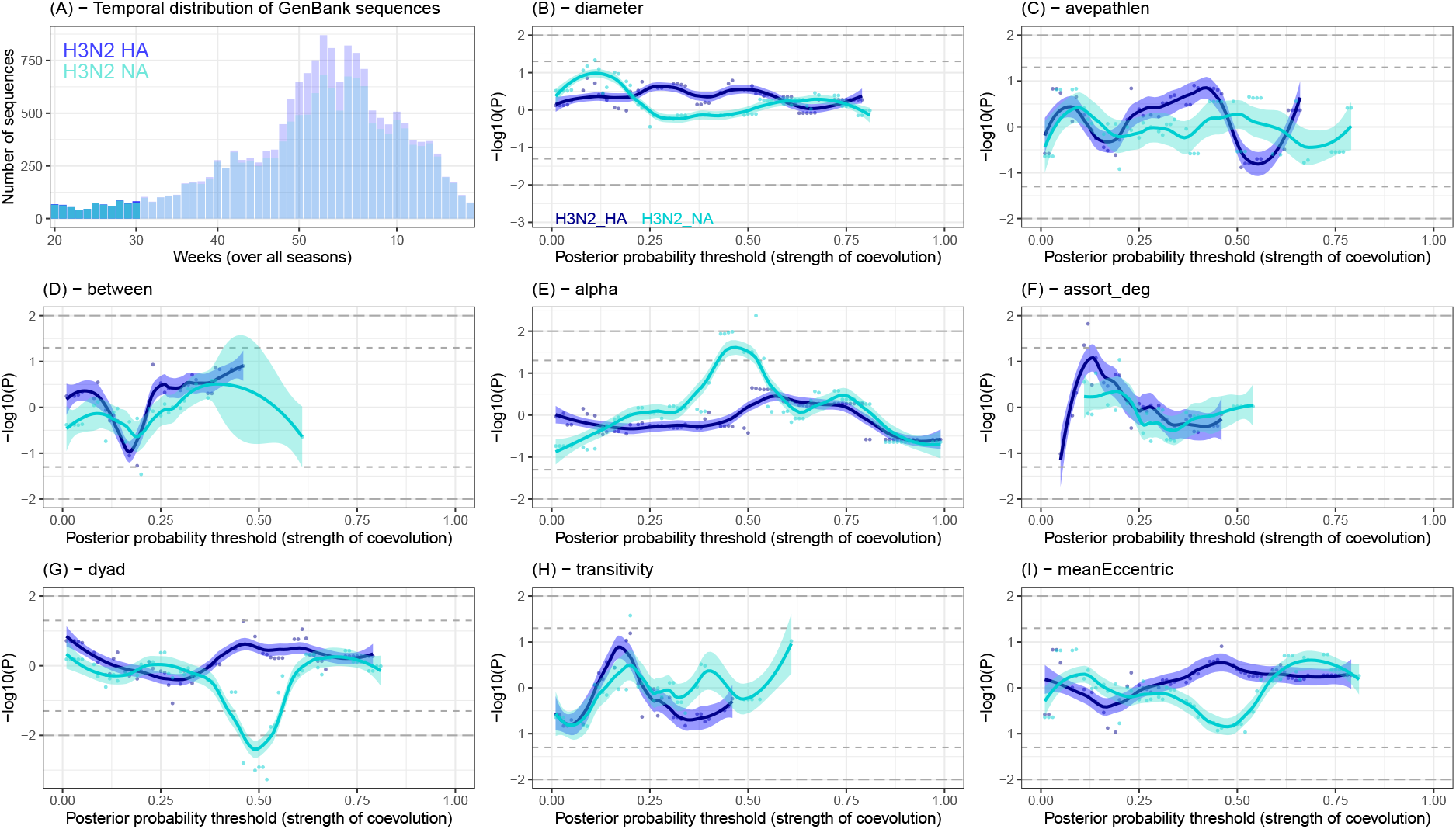
Network analyses based on genetic data from weeks 20-30 (window 1) for the HA and NA genes in H3N2. (A) Temporal distribution of sequences deposited in GenBank (shaded hues for window 4, solid colors for the current window) for HA (blue) and NA (turquoise). The next eight panels show the significance of the robust regressions for each network statistic at a particular posterior probability threshold (the strength of coevolution) against total unweighted ILI value. Negative values indicate a negative slope, and vice-versa for positive values. Gray horizontal lines indicate significance thresholds (dash: 5%; long dash: 1%). LOWESS regressions are shown with the 95% confidence envelope for each subtype and each gene: blue for HA, turquoise for NA.

From window 2 onward, the results are expected to be more meaningful, as they represent a catchment of genetic data directly leading up to the focal season. Indeed, from this point on, HA shows the strongest and most consistent predictive value over a number of network statistics, with temporal trends as we move across windows. More specifically, diameter is first a positive predictor of ILI for HA and NA (Fig. 3B), but as genetic data accumulate, diameter becomes a negative predictor (Fig. 4B and Fig. 5B). This again is highly consistent with our previous results, where during a sever outbreak, networks of coevolving residues become destabilized, and regain stability over time (Aris-Brosou *et al.*, 2017).

**Figure 3.**
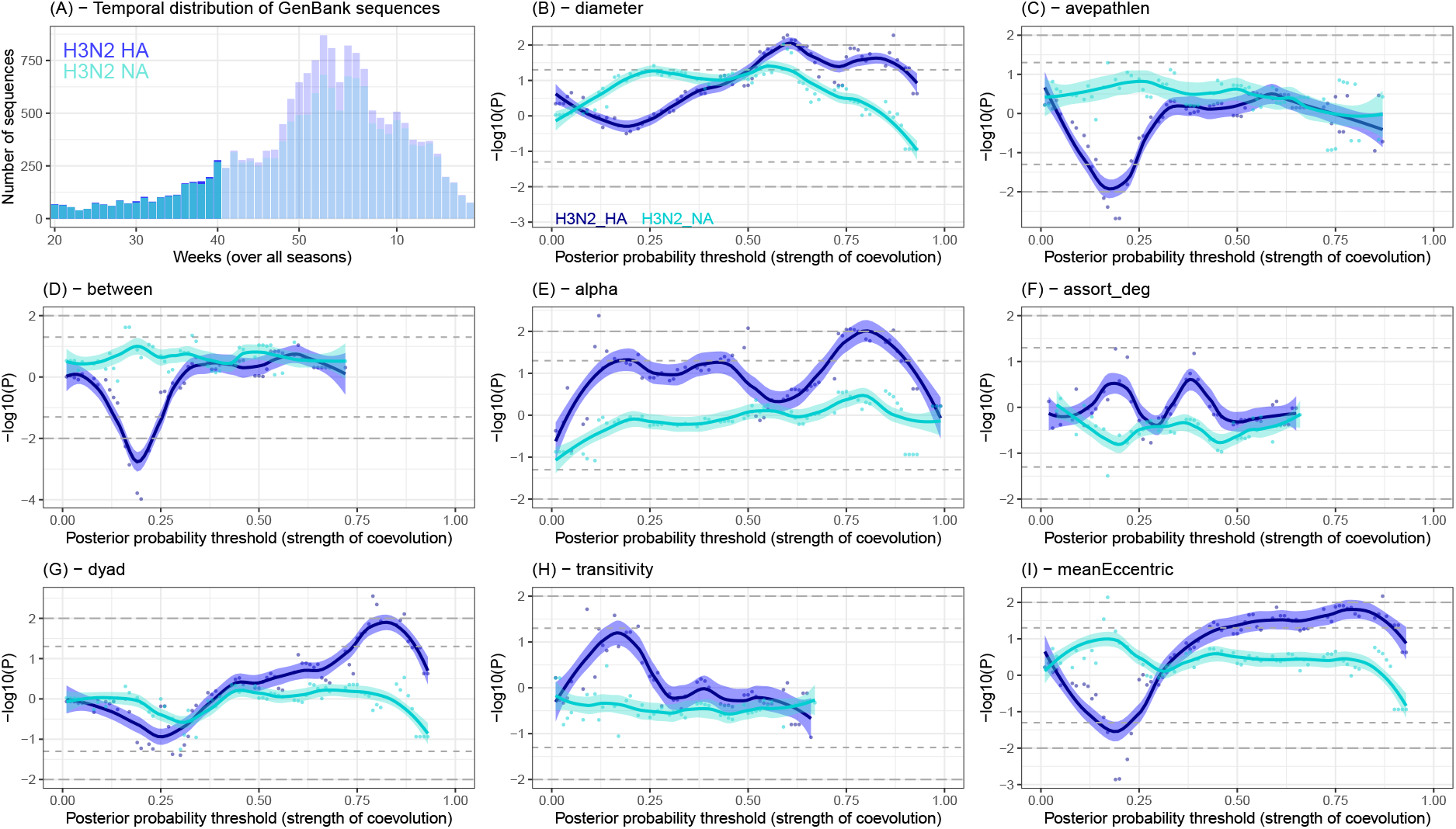
Network analyses based on genetic data from weeks 20-40 (window 2) for the HA and NA genes in H3N2. (A) Temporal distribution of sequences deposited in GenBank (shaded hues for window 4, solid colors for the current window) for HA (blue) and NA (turquoise). The next eight panels show the significance of the robust regressions for each network statistic at a particular posterior probability threshold (the strength of coevolution) against total unweighted ILI value. Negative values indicate a negative slope, and vice-versa for positive values. Gray horizontal lines indicate significance thresholds (dash: 5%; long dash: 1%). LOWESS regressions are shown with the 95% confidence envelope for each subtype and each gene: blue for HA, turquoise for NA.

**Figure 4.**
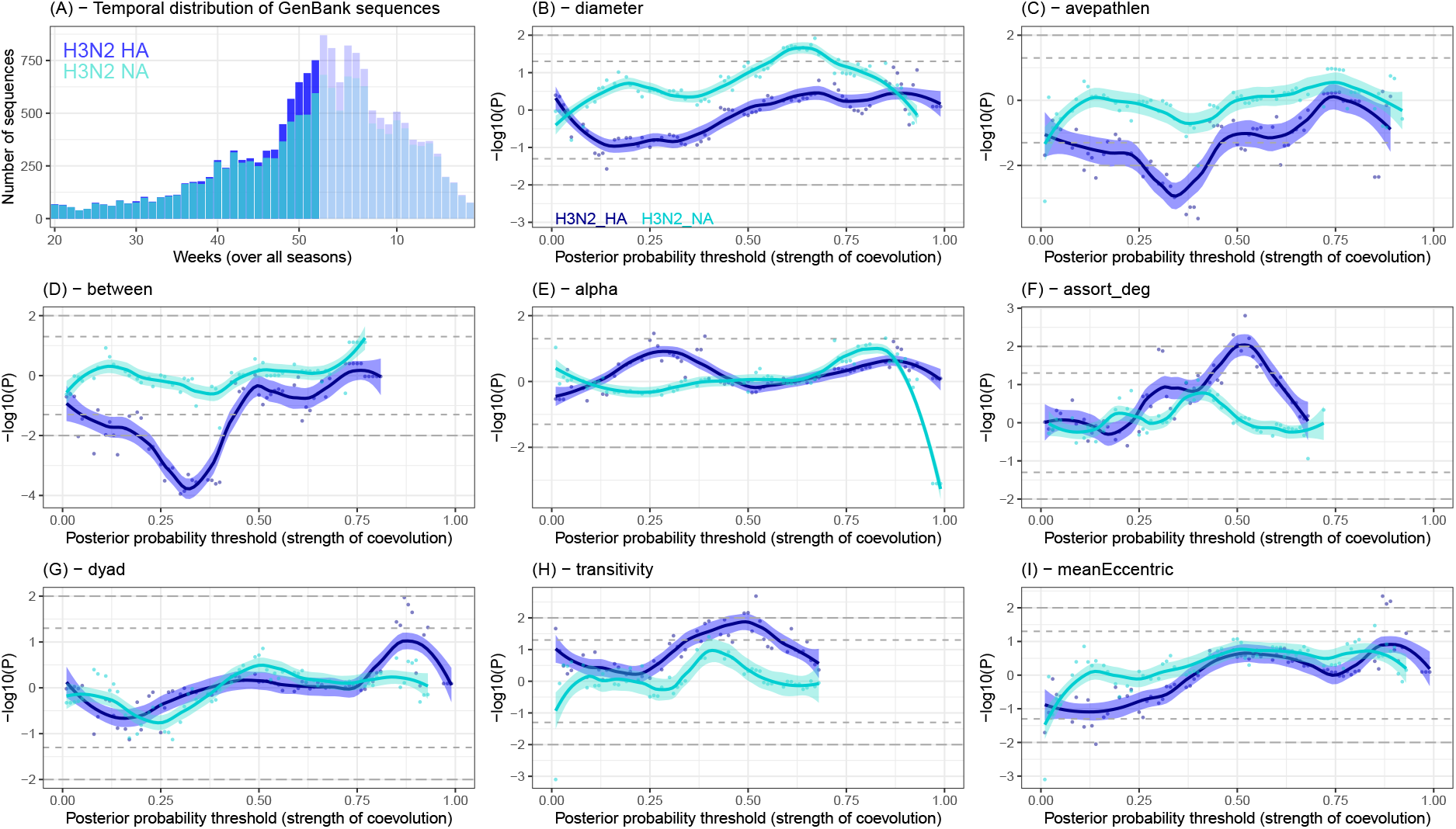
Network analyses based on genetic data from weeks 20-52 (window 3) for the HA and NA genes in H3N2. (A) Temporal distribution of sequences deposited in GenBank (shaded hues for window 4, solid colors for the current window) for HA (blue) and NA (turquoise). The next eight panels show the significance of the robust regressions for each network statistic at a particular posterior probability threshold (the strength of coevolution) against total unweighted ILI value. Negative values indicate a negative slope, and vice-versa for positive values. Gray horizontal lines indicate significance thresholds (dash: 5%; long dash: 1%). LOWESS regressions are shown with the 95% confidence envelope for each subtype and each gene: blue for HA, turquoise for NA.

**Figure 5.**
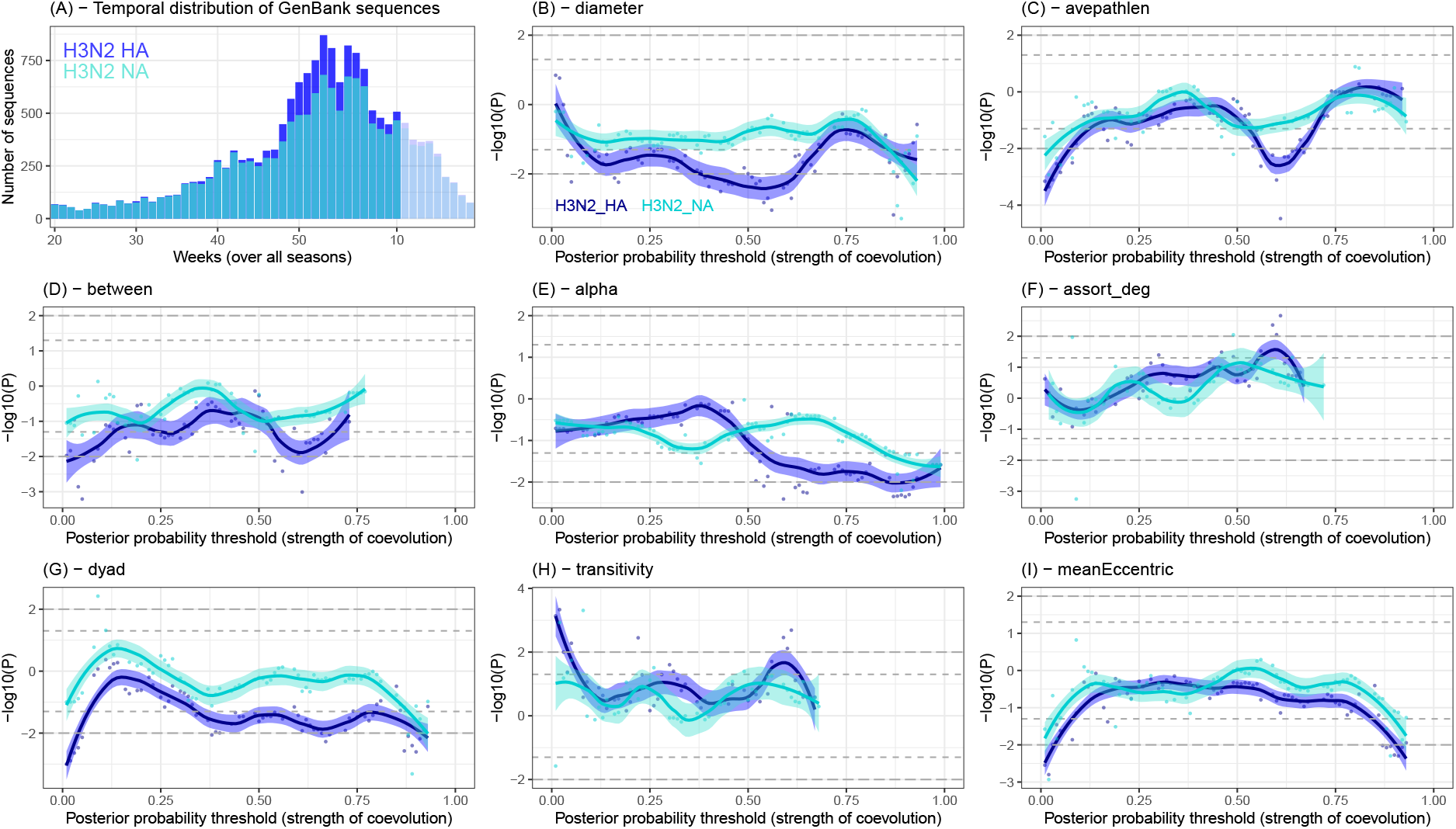
Network analyses based on genetic data from weeks 20-10 (window 4) for the HA and NA genes in H3N2. (A) Temporal distribution of sequences deposited in GenBank (shaded hues for window 4, solid colors for the current window) for HA (blue) and NA (turquoise). The next eight panels show the significance of the robust regressions for each network statistic at a particular posterior probability threshold (the strength of coevolution) against total unweighted ILI value. Negative values indicate a negative slope, and vice-versa for positive values. Gray horizontal lines indicate significance thresholds (dash: 5%; long dash: 1%). LOWESS regressions are shown with the 95% confidence envelope for each subtype and each gene: blue for HA, turquoise for NA.

Likewise, average path length goes from showing no relationship with ILI for sequences sampled sampled 10 weeks prior to the start of the season (Fig. 2C), to progressively showing a more negatively significant trend as we accumulate sequence data through the course of the season, and coevolutionary networks are dislocating. Notably, these networks start by dislocating at weakly coevolving residues (Fig. 3C: *PP* ≈ 0.20), and the stronger interactions are progressively affected (Fig. 4C: *PP* ≈ 0.30; Fig. 5C: *PP* ≈ 0.60). To some extent, the same behavior is observed for betweenness (Fig. 3D: *PP* ≈ 0.20; Fig. 4D: *PP* ≈ 0.30; Fig. 5D: *PP* ≈ 0.60), and alpha centrality (only late in the season: Fig. 5E: *PP* ≥ 0.60). For reasons that are as of yet unexplained, only HA is affected by this trend: NA exhibits no pattern at all.

Similar dynamics are observed for betweenness (from no association to negative to no association for HA: Fig. 2D–Fig. 4D), alpha centrality (from no association to positive to slightly negative for HA: Fig. 2E–Fig. 4E), assortative degree (from no association to positive to no association for HA: Fig. 2F–Fig. 4F), dyad (from no association to briefly positive to no association for HA: Fig. 2G–Fig. 4G), transitivity (from no association to positive to almost no association for HA: Fig. 2H–Fig. 4H), and eccentricity (from no association to positive to no association for HA: Fig. 2I–Fig. 4I). Taken altogether, these results show that networks are destabilized with increasing ILI (smaller diameter) as the epidemic is reaching its peak, and they regain a certain stability at the end of the season, when in-clinic visits plummet. More to the point, these results show that our maximum predictive power is reached when analyzing genetic data collected between week 20 and 40-to-52, that is to say one to fifteen weeks prior to peak ILI burden, which is usually between December and February (Figure 1A; Paget *et al.*, 2007; Murray *et al.*, 2018).

### 3.3 Conclusions

We here showed the existence of an association between the evolutionary dynamics of influenza viruses circulating seasonally in humans and the public health burden caused by this virus in the continental US. More specifically, we showed that network statistics summarizing the level of correlated evolution, and hence evolutionary constraints, affecting the main influenza antigens are associated with the clinical burden due to influenza-like illnesses. With a time-window analysis, we further show that these networks of coevolving residues become destabilized, and regain stability over time (Aris-Brosou *et al.*, 2017).

While the time-window analyses suggest such a seasonal dynamics, causality is far from being obvious. At our level of granularity, it is indeed impossible to identify the drivers of these associations, without also having phenotypic data on virulence and transmissibility of these viruses. Potentially more problematic here is the use and nature of ILI values. By reducing these values to their mean by state by season to represent the entire season, we neglect temporal variation, cumulative ILI values, or maybe more critically maximum ILI values that undoubtedly have a dire impact on health systems. One serious limitation however is that ILI values, irrespective of how they are summarized, do not allow us to tease viral subtypes apart: to alleviate this issue, we limited our analyses to H3N2, assuming that being the prevalent subtype in most seasons (Jester *et al.*, 2020), ILI values will most likely reflect clinic visits due to H3N2, but there is no means of checking the validity of this assumption. Using data that separate ILI by subtype would also be helpful for future analysis. One such database with this capacity would be the WHO/NREVSS (WHO/NREVSS, 2020).

From a purely pragmatic point of view, our results essentially mean that some of these statistics such as network diameter predict ILI values, one to fifteen ahead of peak influenza season in terms of outpatient visits to a clinic. This is a short lead, but one that could henceforth be used routinely to predict ILI burden on the healthcare system, and therefore help plan resource allocations and shifts – and put new non-pharmaceutical interventions into place to curb transmission in the first place (Davies *et al.*, 2020).

## Data availability

The data assembled and the code developed for this work are available from github.com/sarisbro/data.

## Supplementary data

Supplementary data are available at Virus Evolution online.

## Acknowledgements

We thank Compute Canada for providing us with compute time on their servers.

