## Supplementary Information for "Viral evolutionary dynamics predict Influenza-Like-Illnesses in patients"

### Supplementary Figures

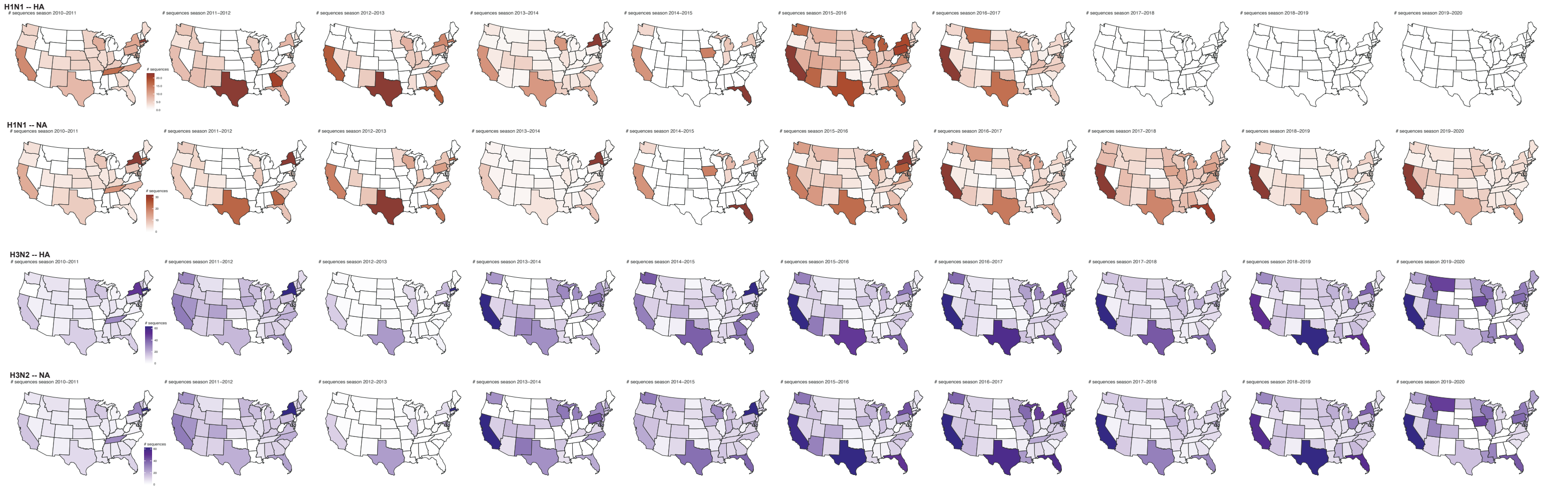


Figure S1: Spatial and temporal distribution of sequence numbers retrieved from GenBank. Numbers are shown for H1N1 HA (top row), NA, (second row), both on a dark red scale, and for H3N2 HA (third row) and NA (bottom row), on a dark blue scale. Note the absence of HA sequences for H1N1 during the last three seasons, 2017-2018 to 2019-2020.

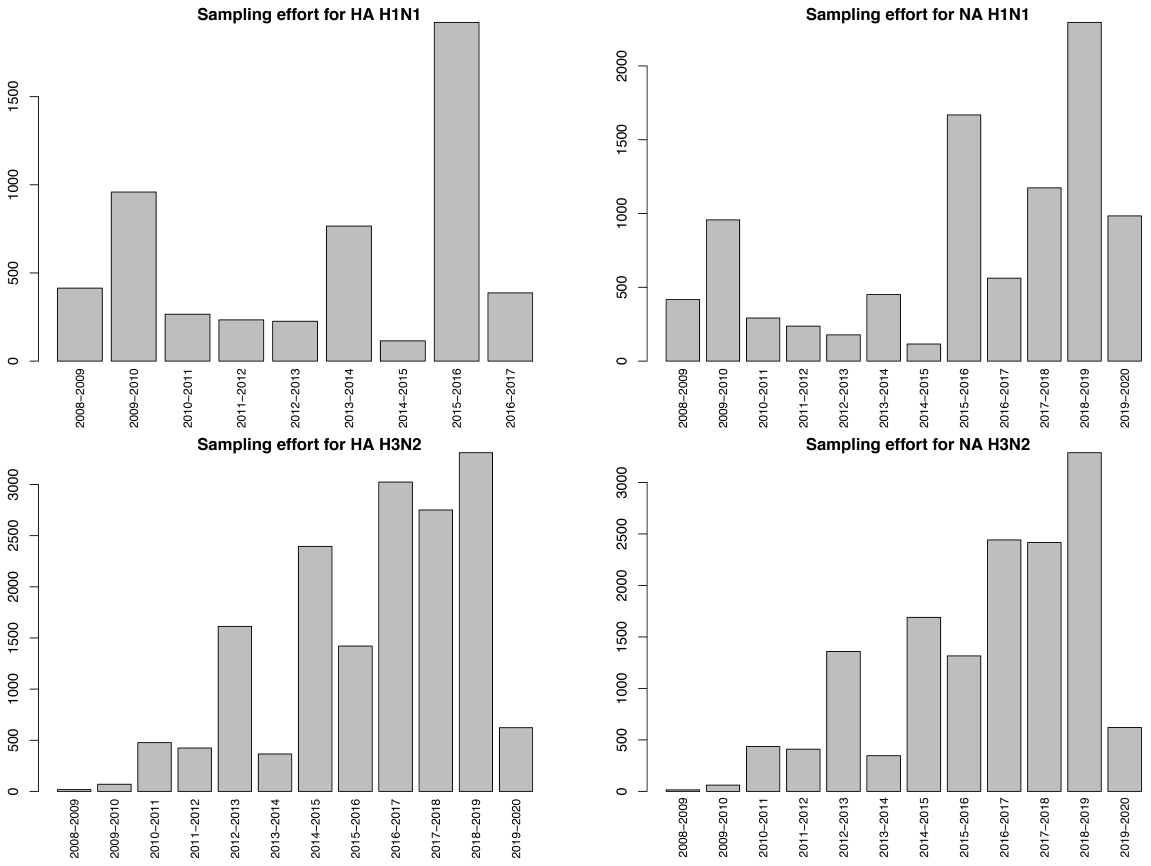


Figure S2: Number of sequences retrieved from GenBank as of April 2020, by season, for H1N1 (top row) and H3N2 (bottom row). Note that the $y$-scales vary across all four panels. Data for the 2008-2009 season are only shown for reference.

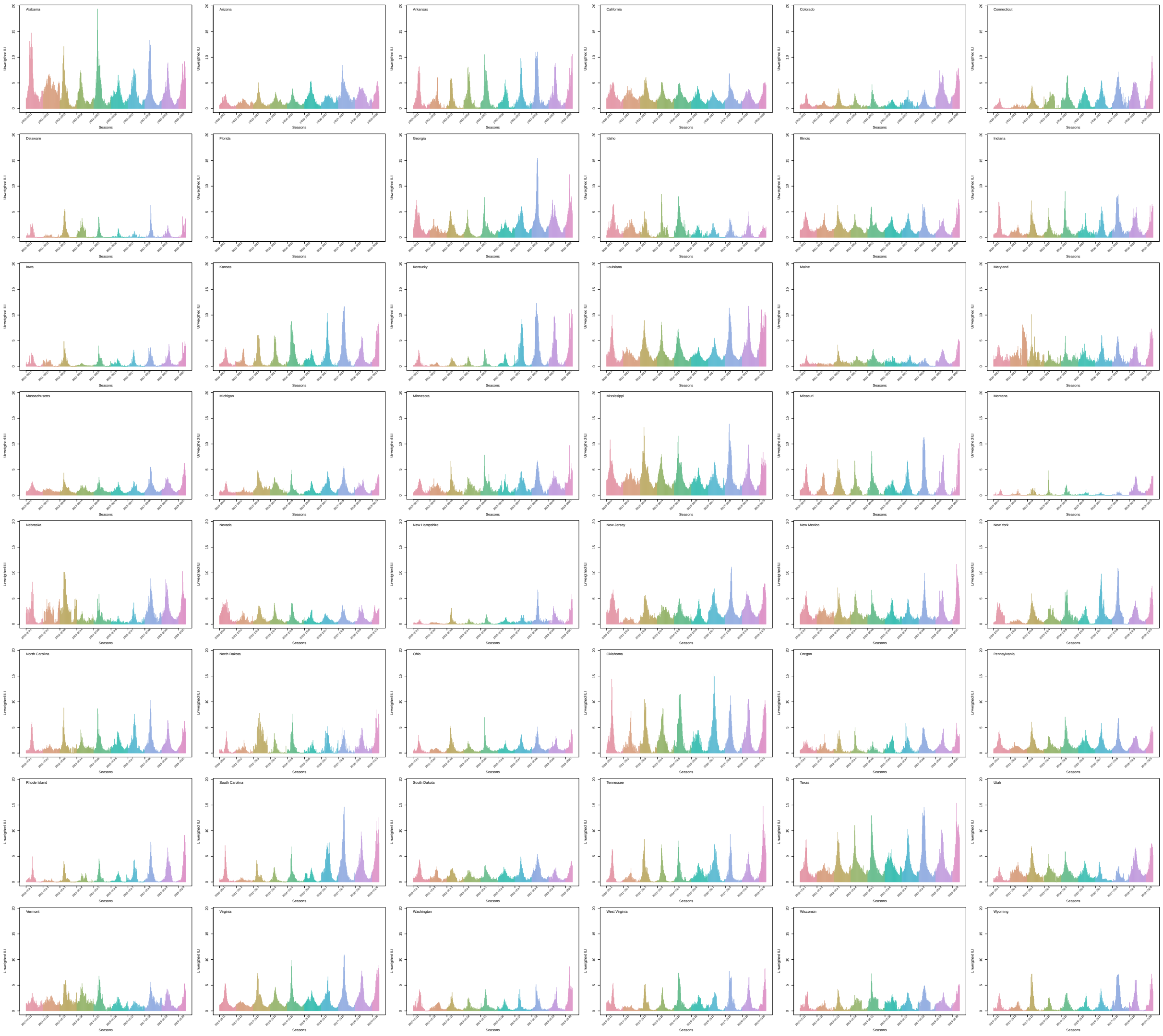


Figure S3: Spatial and temporal distribution of ILI values retrieved from ILI Net. The lower 48 states are ordered alphabetically. Seasons are color coded on a continuous rainbow scale. Note the absence of data for Florida. Note that the scales on the $y$-axes are all identical.

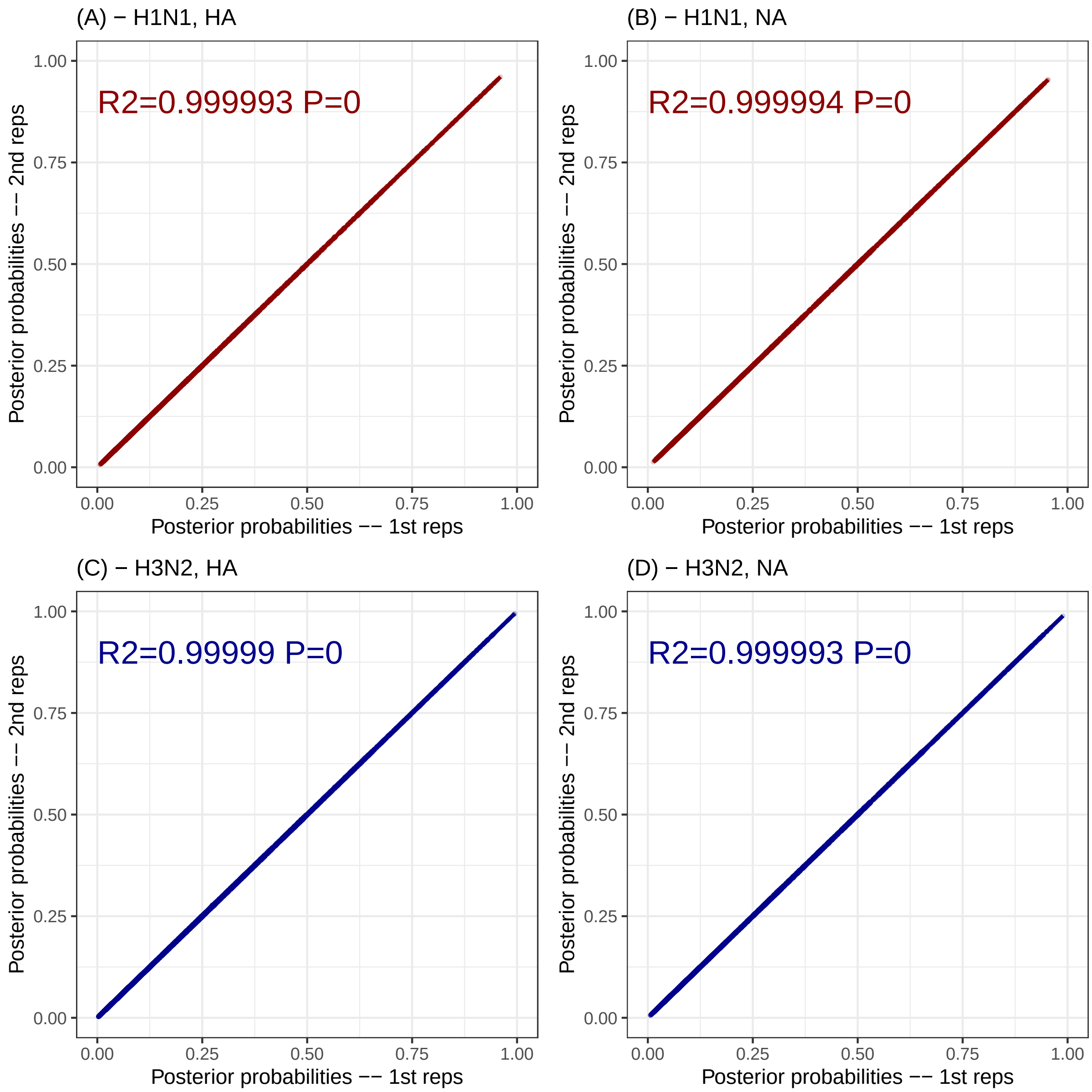


Figure S4: Assessment of the convergence of the Bayesian Graphical Models. Each alignment was analyzed twice, running the complete BGMs and their Markov chain Monte Carlo samplers independently for the H1N1 alignments: (A) HA and (B) NA, and for the H3N2 alignments: (C) HA and (D) NA. Least-square linear models were fit to each dataset in each panel; the $R^2$ is shown, alongside the $P$-value for the estimated slope.

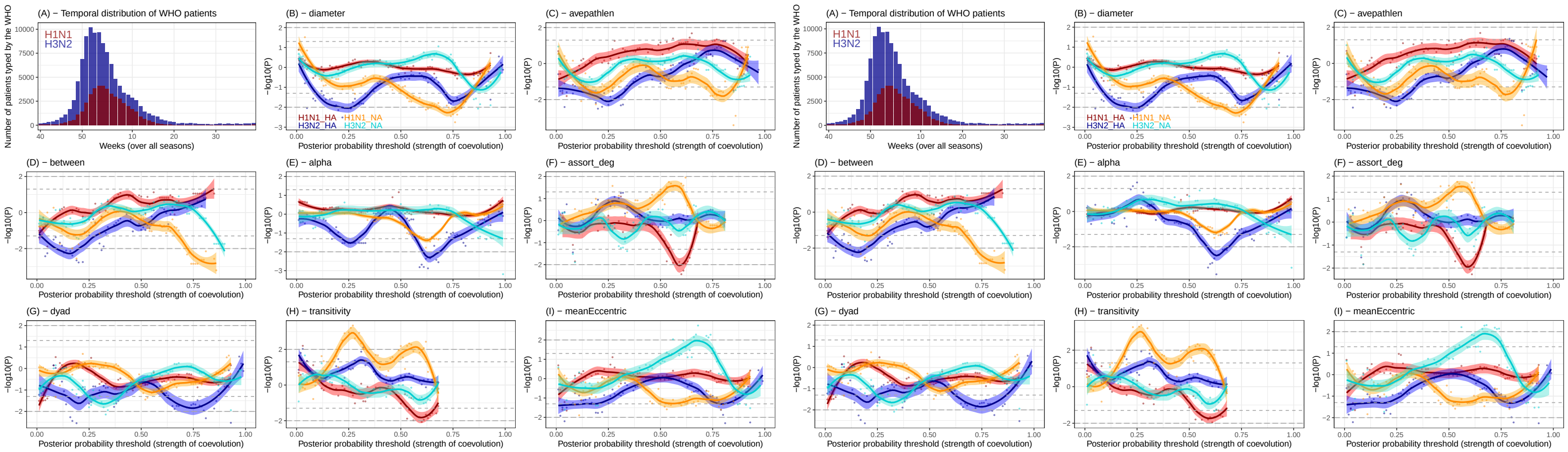


Figure S5: Repeatability of the global analyses for the HA and NA genes of H3N2 and H1N1 subtypes. These global analyses were performed on the whole data, where each alignment was state- and season-specific (from week 40 of a year to week 39 of the following). Run 1 is on the left, run 2 on the right. The distribution of unweighted ILI values summed per week over the entire nine seasons, from 2009-10 to 2018-19 based on WHO data for H1N1 (dark red) and H3N2 (dark blue) is shown on the top left corner for each run. The next eight panels show the significance of the robust regressions for each network statistic at a particular posterior probability threshold (the strength of coevolution) against total unweighted ILI value. Negative values indicate a negative slope, and vice-versa for positive values. Gray horizontal lines indicate significance thresholds (dash: 5%; long dash: 1%). LOWESS regressions are shown with the 95% confidence envelope for each subtype and each gene: warm colors for H1N1 (red for HA, orange for NA) and cold colors for H3N2 (blue for HA, turquoise for NA).

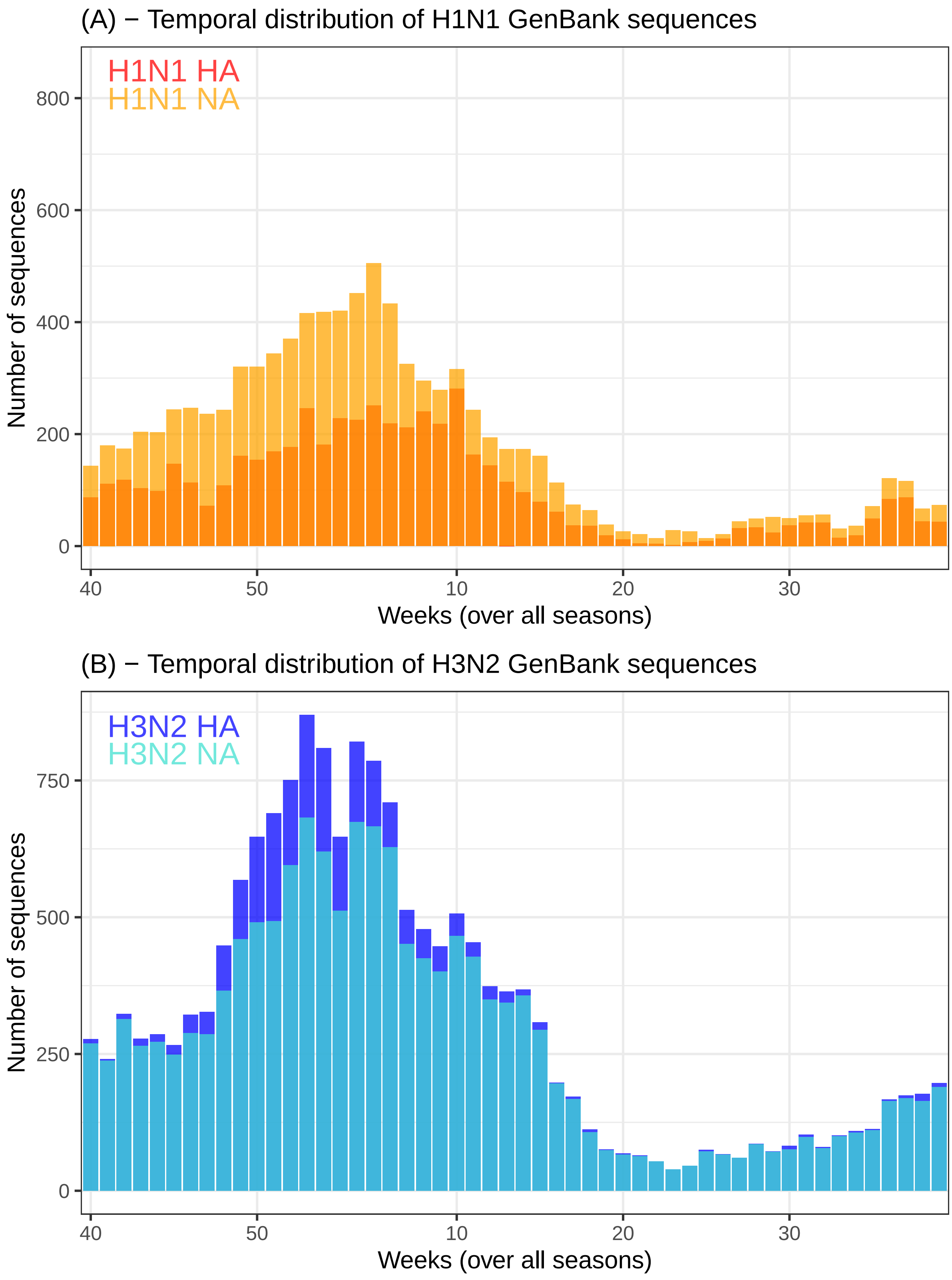




Figure S6: Temporal distribution of sequences retrieved from GenBank aggregated on a weekly basis. (A) H1N1 sequences. (B) H3N2 sequences. Note that the $y$-axes are the same in the two panels.
